## Supplemental Figures for "Sex-dependent gastrointestinal colonization resistance to MRSA is microbiota and Th17 dependent"

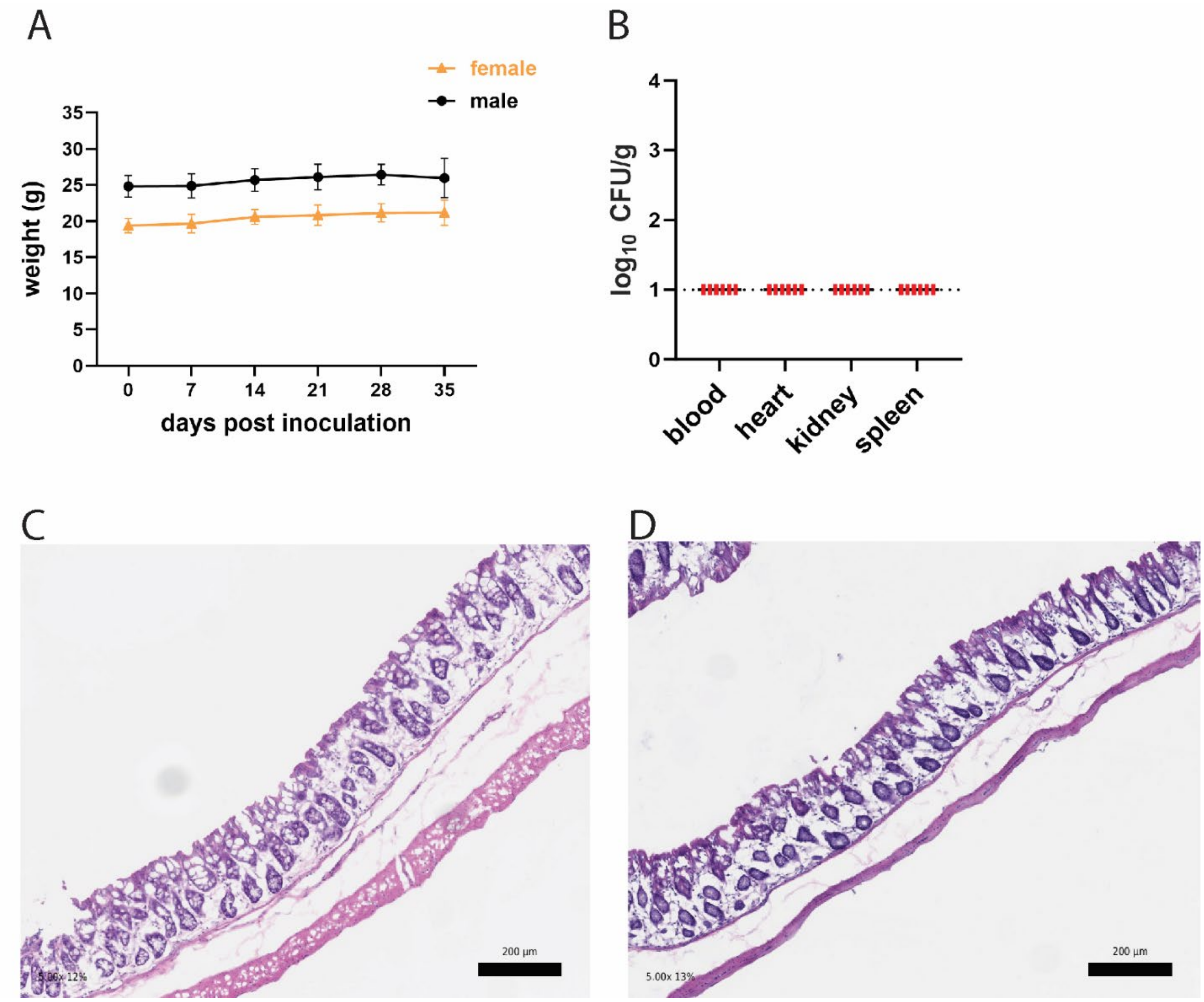


**Supplemental Fig 1. Mice inoculated with MRSA do not display signs of disease.**

**(A)** Weight in grams of B6 mice from NYU and JAX following oral inoculation of MRSA. JAX n=23, NYU n=26. Data points represent mean ± SEM. **(B)** MRSA colony forming units (CFU) in blood, heart, kidney, and spleen following oral inoculation of B6 mice from JAX or NYU. Dots represent individual mice. NYU n=3, JAX n=3. **(C)** Representative image showing hematoxylin and eosin staining of a colonic section 2 days post MRSA inoculation of a male NYU B6 or **(D)** female NYU B6 mouse. 5x magnification, scale bar = 200 µm.

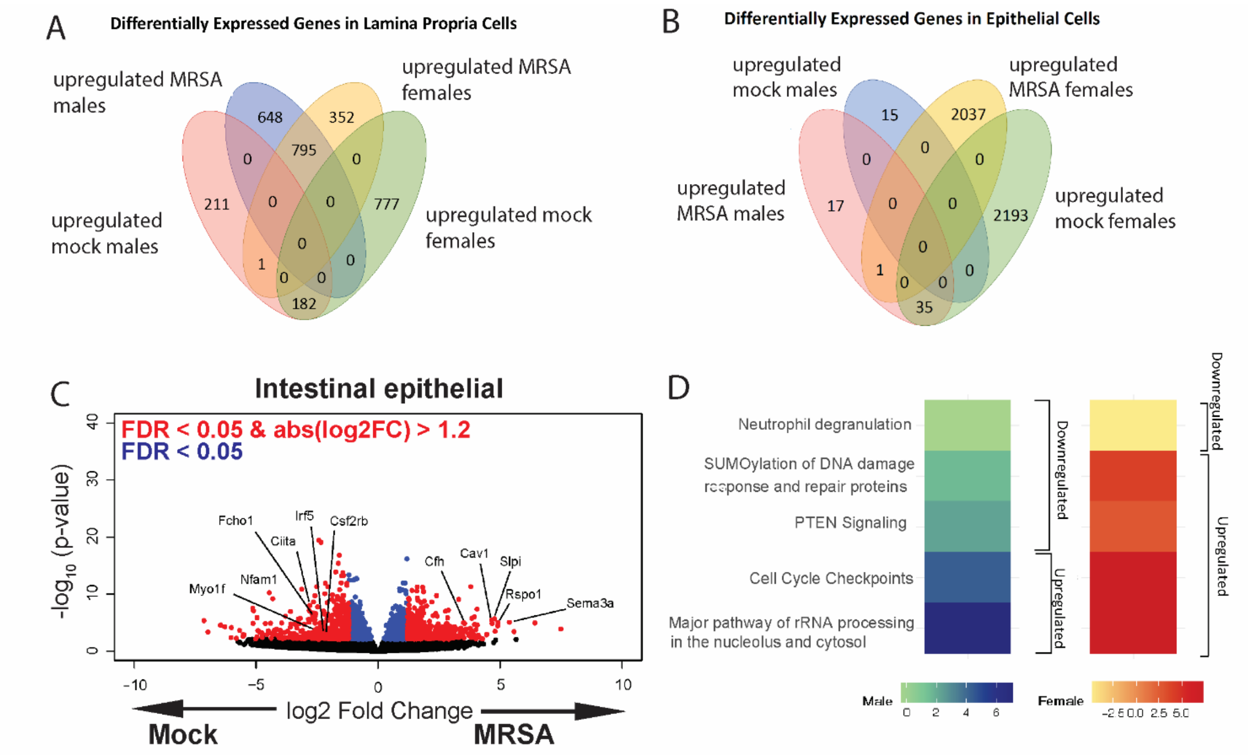


**Supplemental Fig. 2. Transcriptional analyses of the intestine following MRSA inoculation.**

**(A)** Venn diagram showing the number of differentially expressed genes in RNA sequencing (RNA-seq) of lamina propria cells comparing mock to 2dpi MRSA conditions between male and female mice. **(B)** Venn diagram showing the number of differentially expressed genes from RNA seq of intestinal epithelial cells comparing mock and 2dpi MRSA conditions between male and female mice. **(C)** Volcano plot of differentially expressed genes identified by RNA-seq analyses of the intestinal epithelial cells of NYU mice 2 dpi with MRSA compared with PBS mock inoculated NYU mice. **(D)** Ingenuity Pathway Analysis (IPA) of downregulated and upregulated genes in colonic epithelial cells upon MRSA inoculation of male and female mice. Four mice were used for each sequencing experimental condition. Genes shown in red have a false discovery rate (FDR) of <0.05 and an absolute log2 Fold Change (abslog2FC) of >1.2. Genes shown in blue have a FDR of <0.05. Genes linked to X and Y chromosomes were removed from volcano plots.

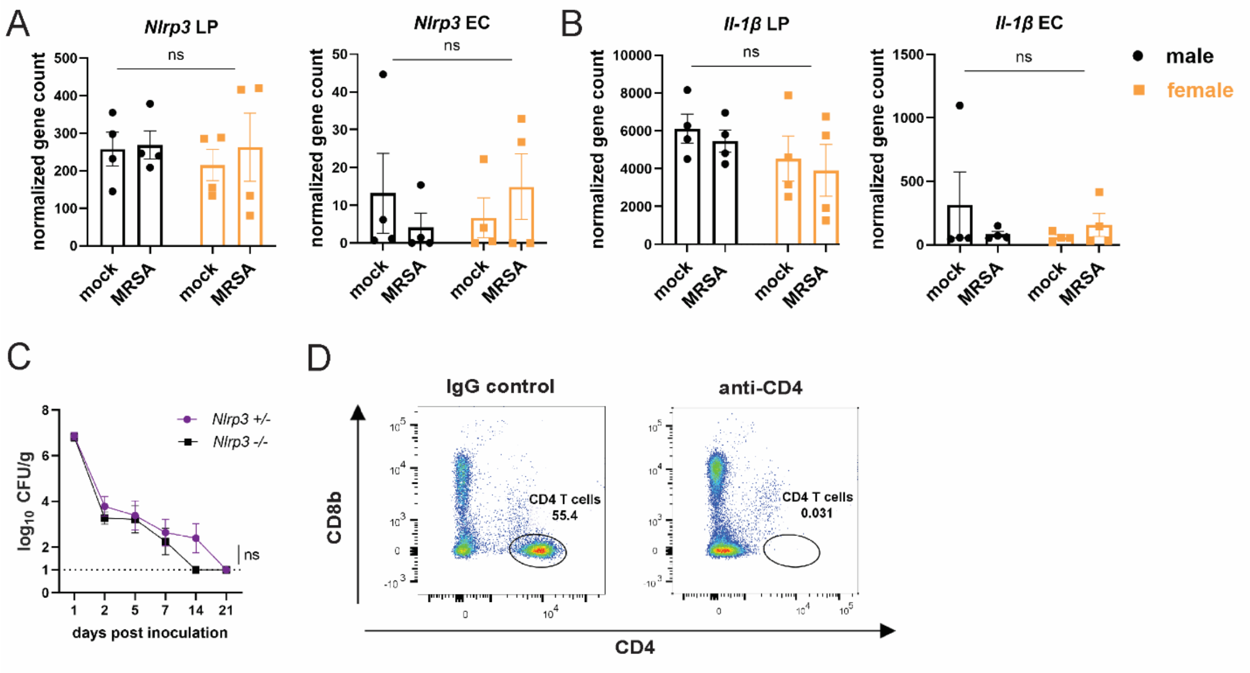


**Supplemental Fig 3. NLRP3 inflammasome activation is not required for MRSA colonization resistance in female NYU mice.**

**(A)** *Nlrp3* gene counts from bulk RNA sequencing (RNA-seq) analysis from cells isolated from the colon lamina propria (LP) or colon epithelial cells (EC). **(B)** *Il-1β* gene counts from bulk RNA seq analysis from cells isolated from the colon lamina propria (LP) or colon epithelial cells (EC). **(C)** MRSA CFU in stool following oral inoculation of *Nlrp3*$^{-/-}$ and *Nlrp3*$^{+/-}$ female mice bred at NYU. *Nlrp3*$^{-/-}$ n=6, *Nlrp3*$^{+/-}$ n=6. **(D)** Representative flow gating to confirm depletion of CD4+ T cells in colon lamina propria 3 days post injection. Data points represent mean ± SEM from at least two independent experiments. Statistical analysis: area under the curve followed by a two-tailed t-test. ns: not significant.

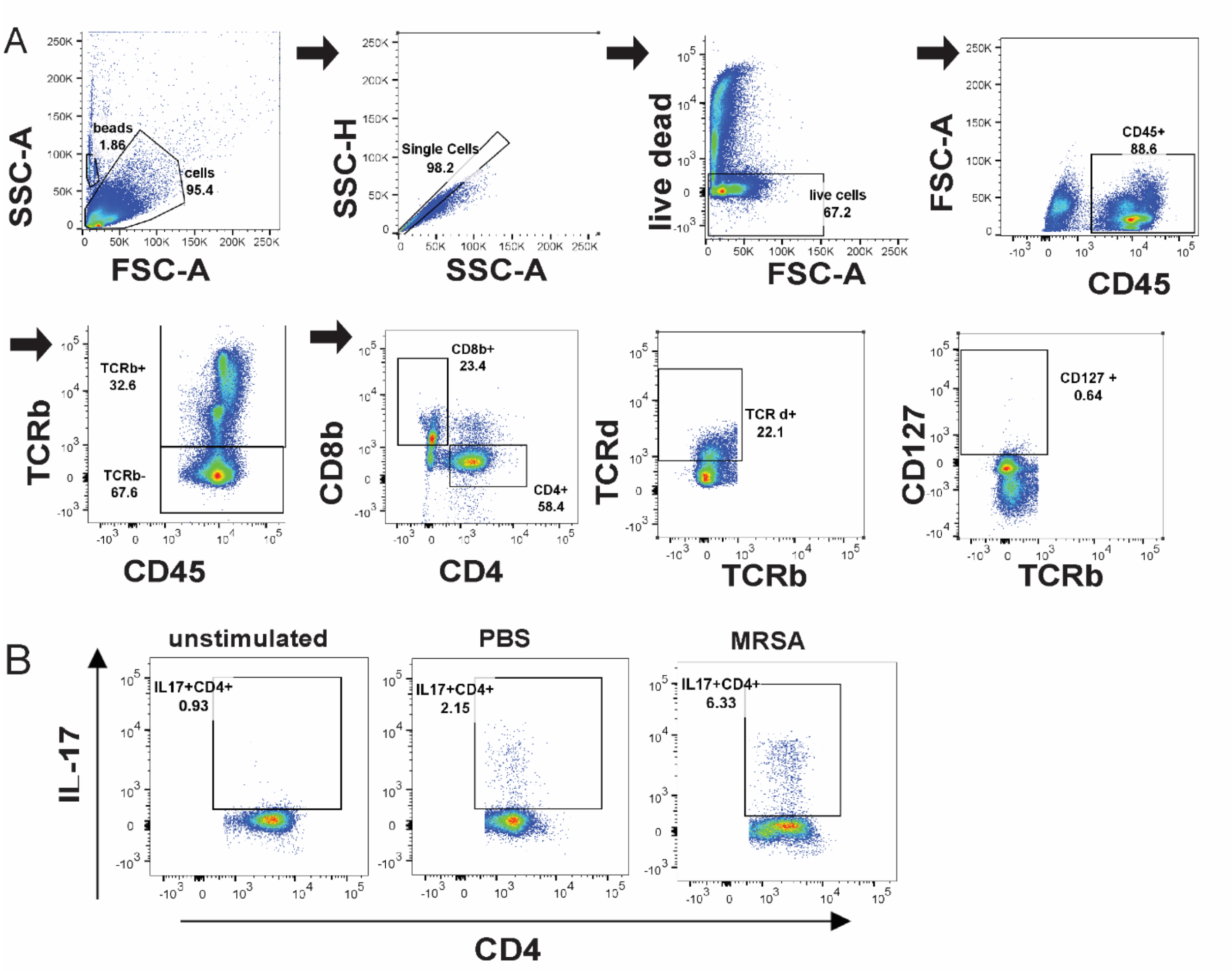


**Supplemental Fig. 4. Flow cytometry gating scheme for lymphoid subsets in the lamina propria.**

**(A)** Flow cytometry gating scheme for ILCs, CD4+ and γδ T cell populations. **(B)** Representative sample gating of IL17A+ CD4+ populations in unstimulated, mock and 2 dpi MRSA treated samples.

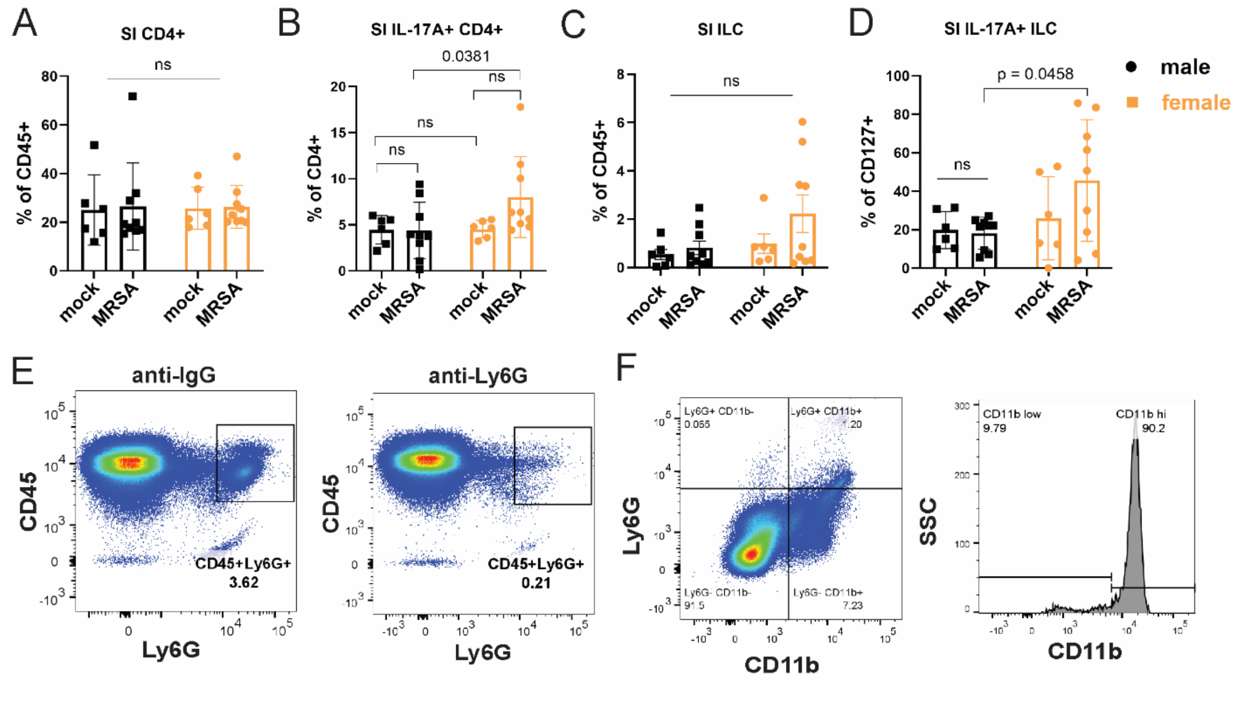


**Supplemental Fig. 5. Immune cell populations detected by flow cytometry.**

**(A)** Flow cytometry of small intestinal (SI) lamina propria CD4+ T cells as a percentage of CD45+ cells in male and female NYU mice treated with PBS or MRSA 2 dpi. **(B)** Flow cytometry of SI lamina propria IL-17A+ CD4+ T cells as a percentage of total CD4+ T cells in male and female NYU mice treated with PBS or MRSA 2 dpi. **(C)** Flow cytometry of cecal-colonic lamina propria CD127+ innate lymphoid cells (ILCs) as a percentage of CD45+ in male and female NYU mice treated with PBS or MRSA 2 dpi. **(D)** Flow cytometry of cecal-colonic lamina propria IL17A+ ILCs as a percentage of total ILCs in male and female NYU mice treated with PBS or MRSA 2 dpi. **(E)** Representative flow gating of CD45+ Ly6G+ cells isolated from the spleens of B6 NYU mice treated with treated with anti-Ly6G neutrophil depleting antibody or anti-IgG control. **(F)** Representative flow cytometry gating plot of CD45+Ly6G+CD11b+ neutrophils isolated from the cecal-colonic tissue of B6 NYU mice treated with a PBS mock control or MRSA. Representative flow cytometry gating plot of CD11b mean fluorescent intensity (MFI) of Ly6G+CD11b+ neutrophils. Data points represent mean ± SEM from at least two independent experiments. Statistical analysis: 2 way ANOVA + Sidak's multiple comparisons test for (A)-(D). ns: not significant.

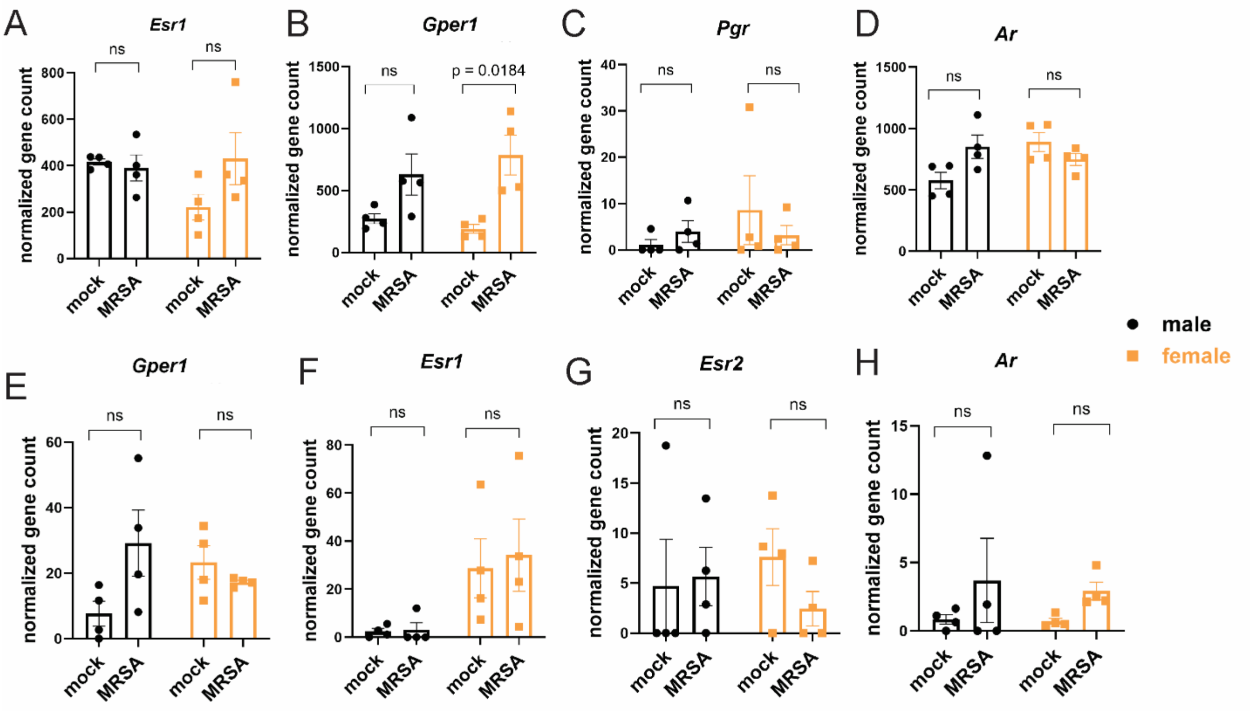


**Supplemental Fig. 6. Sex hormone receptor expression in cecal-colonic lamina propria immune and epithelial cells.**

**(A)** Estrogen receptor alpha (*Esr1*) gene counts **(B)** G-protein coupled estrogen receptor 1 (*Gper1*) expression gene counts **(C)** Progesterone receptor (*Pgr*) gene counts **(D)** Androgen receptor (*Ar*) gene counts. All gene counts are from bulk RNA sequencing analysis from cells isolated from the cecal-colonic lamina propria for **(A) – (D)**. **(E)** G-protein coupled receptor (Gper1) expression gene counts **(F)** Estrogen receptor alpha (Esr1) gene counts **(G)** Estrogen receptor beta (Esr2) gene counts **(H)** Androgen receptor (Ar) gene counts from bulk RNA sequencing analysis. All gene counts are from bulk RNA sequencing analysis from cells isolated from the cecal-colonic epithelial for **(E) – (H)**.  Data points represent mean ± SEM from at least two independent experiments. Statistical analysis: 2 way ANOVA + Sidak's multiple comparisons test. ns: not significant.
